# Occurrence of strawberry viruses in *Fragaria* germplasm and evaluation of cryotherapy as an eradication method for strawberry viruses

**DOI:** 10.1101/2023.07.12.548646

**Authors:** Thomas Wöhner, Monika Höfer

## Abstract

Strawberry plants are highly susceptible to viral infections, which pose significant threats to global strawberry production. This study aims to explore the efficacy of *in vitro* initiation and cryopreservation of shoot tips as a potential strategy for eradicating strawberry viruses. We tested plants for four important strawberry viruses namely: SMoV, SCV, SMEY and SVBV. The plants, which tested positive were either cultivated as in vitro cultures then returned to a green house or field collection cultivation, or treated by cryopreservation. After cryopreservation, the plants were cultivated again in vitro and then in the green house or field. The viruses were detected within each propagation step. Significant eradication effects were found for SMoV and SCV when plants were treated by *in vitro* initiation or with cryotherapy, but not for SMEY or SVBV. The results of this study show that cryotherapy or *in vitro* initiation can lead to the elimination of strawberry viruses, but the kind of therapy appears to depend on the type of virus.

## Text

Strawberries are one of the most economically important temperate fruit crops, with an annual production of 9.157,127.5 t on an area of 389,665 ha worldwide in 2021 (FAO stat, https://www.fao.org/faostat/en). The main producing countries are USA, Netherlands, Morocco, Spain and Albania. With a percentage of 3.1% of the German fruit production, strawberry cultivation was the third-largest in Europe with a yield of 130,630 tonnes on an area of 12,500 ha in 2021 (FAO stat, https://www.fao.org/faostat/en). For the successful cultivation of strawberries, it is necessary to provide virus-free plant material. Virus infections are one main reason for the degeneration of propagation material in strawberries. Once infected, vegetative propagation transmits the viruses from one propagation phase to the next. An infested plant weakens the plant in the long term, leading to increased pathogen susceptibility. However, the virus infection itself also leads to economic losses due to bad fruit quality, deformation of leaves and other symptoms (Martin and Tzanetakis 2006). More than 25 viruses have been described for strawberries to date (Fránová et al. 2019, Koloniuk et al 2022a), which were transmitted via insects, nematodes or other vectors (Bragard et al. 2019, Martin and Tzanetakis et al. 2006, Franova et al. 2019, Koloniuk et al 2022b). Martin and Tzanetakis (2006) reported aphid transmitted viruses, mainly, the *strawberry mottle virus* (SMoV), *strawberry mild yellow edge virus* (SMYEV), *strawberry crinkle virus* (SCV) and *strawberry vein banding virus* (SVBV) as the most economically important ones in strawberry cultivation areas of the world.. Although control of field infestation of the vector *Chaetosiphon fragaefolii* (strawberry aphid) is possible (reviewed in CABI 2022), once a plant is infected, the only way to stop virus dissemination is an eradication of infested plants (Greber 1979, Boxus 1989, Nazarov et al. 2020, Rubio et al. 2020). The generation of virus free plants is an important task for the provision of plants for vegetative propagation, cultivation and preservation of genetic resources. Methods for virus elimination are described for several cultivated plant species and are mainly chemotherapy (Faccioli 2001, Modarresi Chahardehi et al. 2016, AlMaarri et al. 2012), thermotherapy (Faccioli 2001, Wang et al. 2006, AlMaarri et al. 2012, Waswa et al. 2017, Zhao et a. 2018), electrotherapy (AlMaarri et al. 2012), cryotherapy (Zhao et al. 2018) or meristem culture (Faccioli 2001, Quazi and Martin, 1978, Wang et al. 2006, Zhang et al. 2019). For strawberries cryotherapy, thermotherapy and *in vitro* culture techniques were described for single virus eradication (Boxus 1976, McGrew 1965). However, cryotherapy has not been investigated for the eradication of different strawberry viruses. This study investigated the occurrence of strawberry viruses in the germplasm repository in the Fruit Genebank of the Julius Kühn-Institute (JKI) Dresden-Pillnitz and used the well-established method of cryopreservation (Höfer et al. 2016) as a possible method for the eradication of different strawberry viruses.

The plant material was obtained from the *Fragaria* collection of the Fruit Genebank of the Julius Kuehn Institut (JKI). Seventy-seven cultivars and seven unassigned accessions of *Fragaria* ×*ananassa* as well as 168 accession of *Fragaria* wild species and hybrids were tested for four strawberry viruses in the field (see list of the tested cultivars and wild species accession in supplemental material table S1). PCR was used to test and detect four strawberry viruses namely: (SMoV - strawberry mottle virus, SCV – strawberry crinkle virus, SMYEV – strawberry mild yellow edge virus, SVBV – strawberry vain banding virus). A mix of different leaves of up to eight plant samples per accession (n=1-8) were collected (see table S1) for virus detection in the cultivar collection. For initial virus detection in the wild species collection, a mix of different leaves from up to three plants per accession was collected and tested as one sample (n=1). Between three to 10 plant samples (n=3-10) per cultivar were collected for the detection of viruses in the set of 19 cultivars for evaluation of virus eradication efficiency in the field, after in vitro initiation, after cryo-conservation and finally after transfer into the greenhouse again. RNA was isolated from 40 mg leaf material, and the invitrap Spin RNA Mini Kit (Invitec Molecular GmbH, Berlin, Germany) was used for extraction according to the manufacturer’s protocol. The RNA obtained was diluted in 50 μl dd H2O. The quantification of the isolated nucleic acid was performed on the NanoDrop 2000c device. Synthesis of cDNA was performed using the Revert Aid First Strand cDNA Synthesis Kit (Thermo Fisher) according to the manufacturer’s protocol. Random hexamer oligos and oligo_dT18-nucleotides were used for the synthesis. A total of 1 μg RNA was the input amount for cDNA synthesis per sample. Successful cDNA synthesis was evaluated using a standard PCR method using elongation factor EF specific primers (EF_F und EF_R, Flachowsky et al. 2007). The PCR conditions were: 13,4 μl ddH2O, 2.5 μl 10x DreamTaq Puffer with 20 mM MgCl2, 2.5 μl dNTPs with 2 mM, 1.25 μl of 10 μM EF1αF and EF1αR, 1 μl 20x red buffer, 1 μl of 0,125% BSA (Zhang et al. 2014), 1 μl of 25% PVP (Koonjul et al. 1999) and 0,1 μl of 5 U μl-1 DreamTaq polymerase. A total of 1μl cDNA was used for each PCR reaction. PCR was performed with 1 x initial denaturation: 94 °C 3’, 35 x denaturation/annealing/elongation: 94 °C 30’’/56 °C 1’/72 °C 1’, 1 x final elongation: 72 °C 3’ and 1 x cooling: 10 °C ∞. The primer sequences to proof strawberry leaf material on the occurrence of strawberry viruses was obtained from the publication listed in table S2 and PCR was performed according to the mastermix and conditions in table S3. Amplificates of investigated samples, positive and negative control samples will be separated by agarose gel electrophoresis. For each sample 10 μl PCR product is loaded into a 1,5 % agarose gel and separated at 90 Volt. A 50 bp size standard (Thermo-Fisher Scientific) is used. Positive samples amplify the specific fragment, whereas negative samples obtained no fragment. The evaluation of virus eradication effect by cryotherapy compared to *in vitro* initiation was tested on 19 cultivars (Coral, Dukat, Florika, Fraginetta, Gloria, Mieze Nova, Mrak, Pantagruella, Papa Lange, Pegasus, Pervagata, Polka, Rosa Perle, Rubia, Senga Dulcita, Senga Gigana, Symphony, Talisman, Triscana). For the evaluation of virus eradication effect by cryotherapy, samples of the cultivars were obtained from the field (test phase – A). Stolons of positive tested plants were obtained and shoot tips were isolated in the laboratory according to the experimental procedures described in Höfer (2011). Up to three single shoot tips of virus positive plants (n= up to 3) were dissected to establish *in vitro* cultures before cryotherapy (test phase – B). Negatively tested plants obtained from *in vitro* cultures were used for re-transmission from the laboratory into the greenhouse (test phase – C) for virus retesting to study the effect of shoot tip dissection on virus elimination. Up to three individual plants (n= up to 3) were used for virus testing. *In vitro* apical shoot tips from positive tested cultivars were dissected from up to 4-week-old *in vitro* plants and the method described in Höfer et al. 2016 was performed for cryopreservation and recovery of plant shoot tips. Up to 10 plants per cultivar were (n= up to 10) were tested on the occurrence of viruses (in-vitro culture after cryo, test phase – D). After transmission of recovered plants into the greenhouse (test phase – E, plants were tested on the occurrence of viruses as described for initial virus testing (n= up to 9). The frequency of positive tested plant samples per virus, the percentage of positive and negative tested cultivars/species was calculated. For the 19 cultivars mentioned above, the frequency of positive tested samples for each virus was calculated per cultivar and test phase.

A total of 84 *Fragaria* ×*ananassa* accessions and 164 accessions of 22 *Fragaria* wild species and hybrids were tested on the occurrence of four strawberry viruses. An example of the detection results obtained by PCR for each single virus is shown in figure 1. Table 1 shows the percentage of positive tested samples. The highest virus frequency in *Fragaria ×ananassa* was observed for the SCV (73.2%) and SMYEV (72.1%). A lower frequency was obtained for SMoV (57.5%) and SVBV (4.3%). Single virus frequencies determined for each species are shown in table 1. To determine the frequency of each virus over all tested samples and species, a mean frequency of each virus was calculated. The most frequent virus was SMYEV (74.5%), whereas SCV (35.9%), SMoV (30.9%) and SVBV (11.8%) showed a lower mean frequency. Between 6.8 % (SVBV) and 82.9 % (SMYEV) of samples collected from 19 cultivars tested positive for all four viruses in field. The effect of virus elimination when shoot tips were isolated from stolons of infected strawberry plants to establish *in vitro* cultures resulted in 26.3 % (SMYEV) to 98.2 % (SVBV) virus free plants. After retransmission into the green house between 2.6 % (SCV) to 76.3 % (SMYEV) of the tested plants were re-infected with viruses (table 2). The effect of cryotherapy was also investigated and 14.9 % (SMYEV) to 100 % (SCV) negative tested plants were obtained. After re-transmission of cryotherapy threated plants into the green house between 22.6 % (SMYEV) and 100 % (SCV) of plants tested negative on the strawberry viruses. The results are shown in table 3.

**Table 1.** Frequency of four strawberry viruses in *Fragaria* germplasm.

| Species | No. of samples | % positive tested samples* / accessions |  |  |  |
| --- | --- | --- | --- | --- | --- |
|  |  | SMoV | SMYEV | SCV | SVBV |
| *Fragaria ×ananassa* | 280 | 57.5 | 72.1 | 73.2 | 4.3 |
| *Fragaria ×bifera* | 3 | 100.0 | 100.0 | 100.0 | 66.7 |
| *Fragaria ×bringhurstii* | 1 | 0.0 | 100.0 | 100.0 | 0.0 |
| *Fragaria bucharica* | 8 | 12.5 | 75.0 | 37.5 | 0.0 |
| *Fragaria chiloensis* | 19 | 36.8 | 100.0 | 78.9 | 15.8 |
| *Fragaria corymbosa* | 9 | 22.2 | 44.4 | 0.0 | 33.3 |
| *Fragaria gracilis* | 4 | 50.0 | 0.0 | 0.0 | 0.0 |
| *Fragaria hybr.* | 2 | 50.0 | 100.0 | 50.0 | 0.0 |
| *Fragaria iinumae* | 1 | 0.0 | 100.0 | 0.0 | 100.0 |
| *Fragaria mandshurica* | 8 | 25.0 | 75.0 | 25.0 | 0.0 |
| *Fragaria moschata* | 10 | 30.0 | 10.0 | 40.0 | 10.0 |
| *Fragaria moupinensis* | 1 | 0.0 | 0.0 | 0.0 | 0.0 |
| *Fragaria nilgerrensis* | 5 | 0.0 | 100.0 | 0.0 | 20.0 |
| *Fragaria nipponica* | 7 | 14.3 | 100.0 | 42.9 | 0.0 |
| *Fragaria nubicola* | 6 | 0.0 | 100.0 | 33.3 | 0.0 |
| *Fragaria orientalis* | 7 | 28.6 | 100.0 | 28.6 | 0.0 |
| *Fragaria pentaphylla* | 3 | 66.7 | 0.0 | 0.0 | 0.0 |
| *Fragaria sp.* | 19 | 5.3 | 42.1 | 5.3 | 0.0 |
| *Fragaria tibetica* | 4 | 0.0 | 100.0 | 0.0 | 0.0 |
| *Fragaria vesca* | 18 | 38.9 | 100.0 | 83.3 | 5.6 |
| *Fragaria virginiana* | 16 | 43.8 | 93.8 | 50.0 | 0.0 |
| *Fragaria viridis* | 6 | 100.0 | 100.0 | 50.0 | 16.7 |
| *Fragaria yezoensis* | 7 | 28.6 | 100.0 | 28.6 | 0.0 |

\* in case of *Fragaria ×ananassa* more then one sample per accession were tested

**Table 2.** Results from the evaluation of strawberry virus eradication via stolon meristem explant isolation and re-transmission into the green house

| No. | test phase | plant material | virus | samples tested | no. positive | no. negative | %- pos. | %-difference from 100 % pos. tested plant material |
| --- | --- | --- | --- | --- | --- | --- | --- | --- |
| A | from field collection | leaves from flower box | SMoV | 76 | 51 | 25 | 69.8 | - |
|  |  |  | SMYEV |  | 58 | 18 | 82.9 | - |
|  |  |  | SCV |  | 58 | 18 | 73.3 | - |
|  |  |  | SVBV |  | 5 | 71 | 6.8 | - |
| ↓ only positive plants from phase (I) were used in phase (II) |  |  |  |  |  |  |  |  |
| B | in-vitro culture before cryo | explants from stolon meristem | SMoV | 47 | 15 | 32 | 31.6 | 68.4 |
|  |  |  | SMYEV |  | 35 | 12 | 73.7 | 26.3 |
|  |  |  | SCV |  | 3 | 44 | 6.1 | 93.9 |
|  |  |  | SVBV |  | 1 | 46 | 1.8 | 98.2 |
| ↓ only negative plants from phase (II) were used in phase (III) |  |  |  |  |  |  |  |  |
| C | re-transmission from the lab into green house | leaves from flower box | SMoV | 49 | 8 | 41 | 17.5 | 82.5 |
|  |  |  | SMYEV |  | 38 | 11 | 76.3 | 23.7 |
|  |  |  | SCV |  | 2 | 47 | 2.6 | 97.4 |
|  |  |  | SVBV |  | 2 | 47 | 4.4 | 95.6 |

**Table 3.** Results from the evaluation of strawberry virus eradication via stolon meristem explant isolation, cryopreservation treatment and re-transmission into the green house

| No. | test phase | plant material | virus | samples tested | no. positive | no. negative | %-pos. | %-difference from 100 % pos. tested plant material |
| --- | --- | --- | --- | --- | --- | --- | --- | --- |
| ↓ only positive plants from phase (I) were used in phase (II) |  |  |  |  |  |  |  |  |
| B | in-vitro culture before cryo | explants from stolon meristem | SMoV | 47 | 15 | 32 | 31.6 | 68.4 |
|  |  |  | SMYEV |  | 35 | 12 | 73.7 | 26.7 |
|  |  |  | SCV |  | 3 | 44 | 6.1 | 93.9 |
|  |  |  | SVBV |  | 1 | 46 | 1.8 | 98.2 |
| ↓ only positive plants from phase (II) were used in phase (III) |  |  |  |  |  |  |  |  |
| D | in-vitro culture after cryo | explants from stolon meristem | SMoV | 111 | 12 | 99 | 9.0 | 91.0 |
|  |  |  | SMYEV |  | 99 | 12 | 85.1 | 14.9 |
|  |  |  | SCV |  | 0 | 111 | 0 | 100.0 |
|  |  |  | SVBV |  | 2 | 109 | 1.1 | 98.9 |
| ↓ only negative plants from phase (III) were used in phase (IV) |  |  |  |  |  |  |  |  |
| E | re-transmission from the lab after cryo | leaves from flower box | SMoV | 108 | 10 | 98 | 9.1 | 90.9 |
|  |  |  | SMYEV |  | 84 | 24 | 77.4 | 22.6 |
|  |  |  | SCV |  | 0 | 108 | 0 | 100.0 |
|  |  |  | SVBV |  | 1 | 107 | 0.4 | 99.6 |

**Figure 1.**
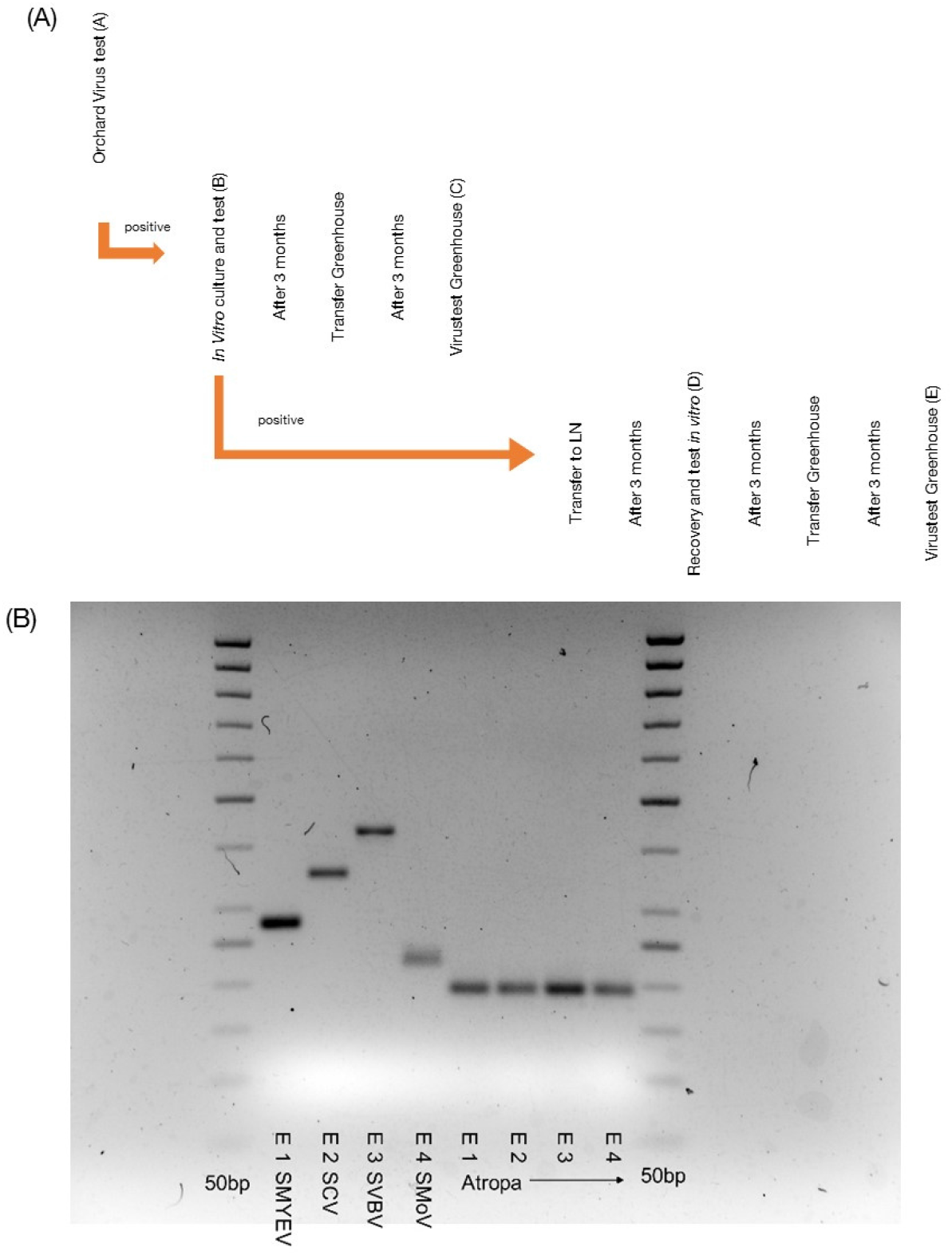
(A) Schematic phases of virus elimination by in vitro initiation and cultivation or cryopreservation (A-E). (B) Detection of four strawberry viruses with RT-PCR. E1 SMYEV –positive control of strawberry mild yellow edge virus with a band at 271 bp, E2 SCV –positive control of strawberry crincle virus with a band at 345 bp, E3 SVBV –positive control of strawberry vein banding virus with a band at 435 bp, E4 –positive control of strawberry mottle virus with a band at 219 bp, E1-E4 –negative control using the AtropaNad2 band at 188 bp, 50bp – size marker.

Strawberries are highly susceptible to strawberry viruses, and sources of resistance to the viruses or vectors are not investigated so far (Shanks and Barrit 1974, Barrit and Shanks 1980). Chemical controls against the vectors are also possible, but only with very high application rates, which is contrary to current socio-economic developments. Once a plant is infected, it can only be eradicated and new virus-free plant material has to be made available. Providing virus-free plant material for new plantings is therefore the best strategy so far (Bettoni et al. 2022). In this study, we therefore investigated the effect of *in vitro* initiation and cryopreservation on virus elimination on strawberry (Figure 1A). Significant eradication effects were found for all viruses by *in vitro* initiation and further by cryopreservation (Table 2 and 3). In potatoes, Bettonie et al. 2022 and Kushnarenko et al. 2017 showed a high elimination rate against three viruses by chemotherapy and cryotherapy. In other species such as raspberry (35%), sweet potato (100%), banana (34 to 90%), grapevine (96% to

100%), quince (33 to 37%), apple (35 to 100%) and *Prunus* spec. (50%), cryotherapy was also successfully performed to eliminate viruses (Harding et al. 2004, Helliot et al. 2002, Feng et al. 2013, Cui et al. 2015; Pathirana et al. 2015, 2019, Farhadi-Tooli et al. 2022, Wang et al. 2022a). In addition to cryotherapy, this study confirms that *in vitro* initiation (Table 2) already leads to a reduction on strawberry viruses, which was previously shown by Boxus (1976). However, the experiments also show that no effect could be detected for the eradication of SMYEV. Binhua et al. (2008) especially reports the successful elimination of SMYEV by freezing, which is contradictory to the results of that study. This virus showed the highest frequency in the tested plant assortment. Whether this virus can be successfully eliminated in combination with heat or chemotherapy (Bettonie et al. 2022) remains to be answered in future research projects.

## Acknowledgements

We would like to thank Sabine Bartsch for her technical assistance with sampling, DNA isolation and virus testing, and Ute Sonntag and Katrin Winkler for her assistance with inculturing and cryotherapy of strawberries.

## Supplemental tables

**Table S1.**
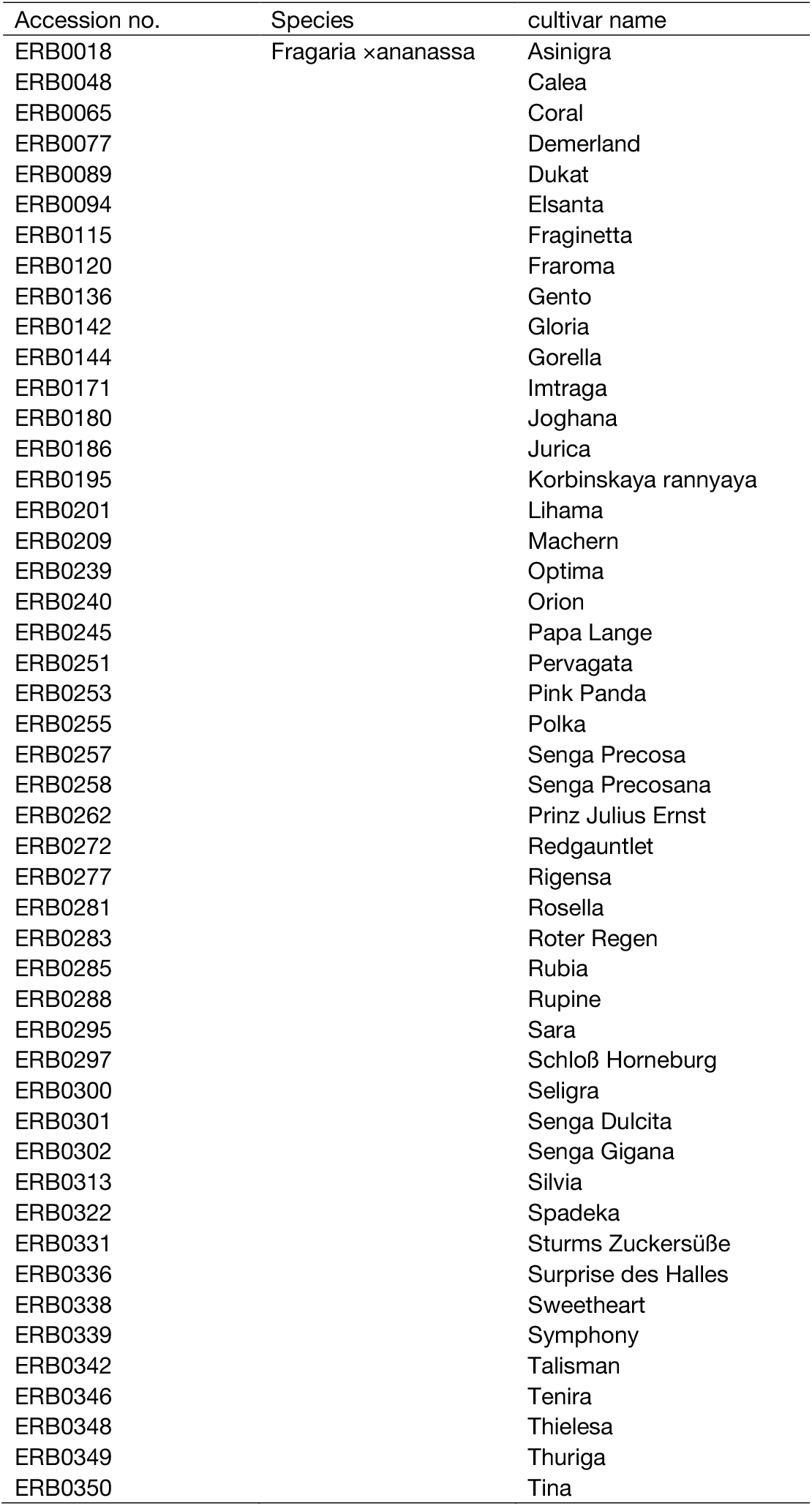
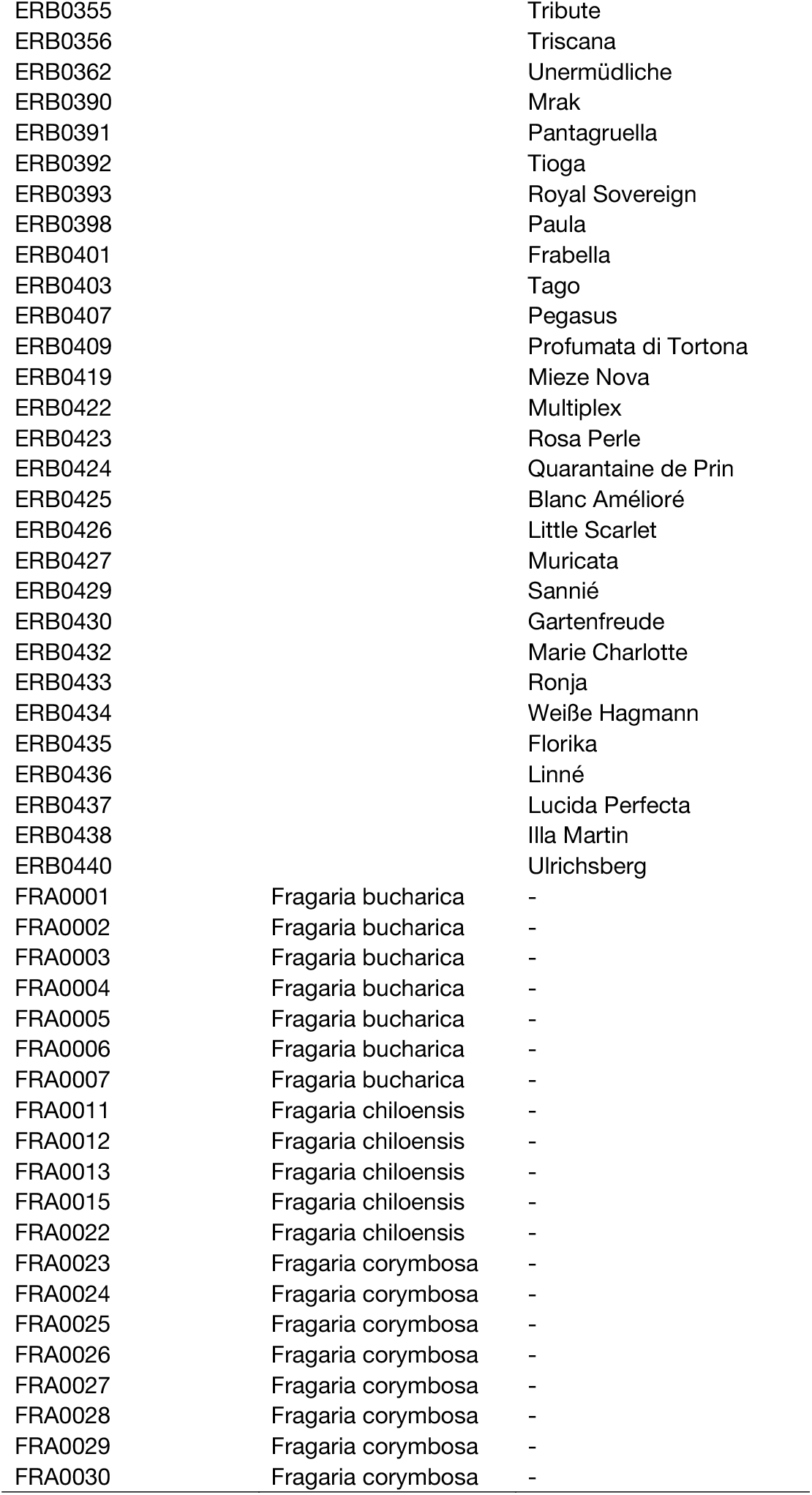
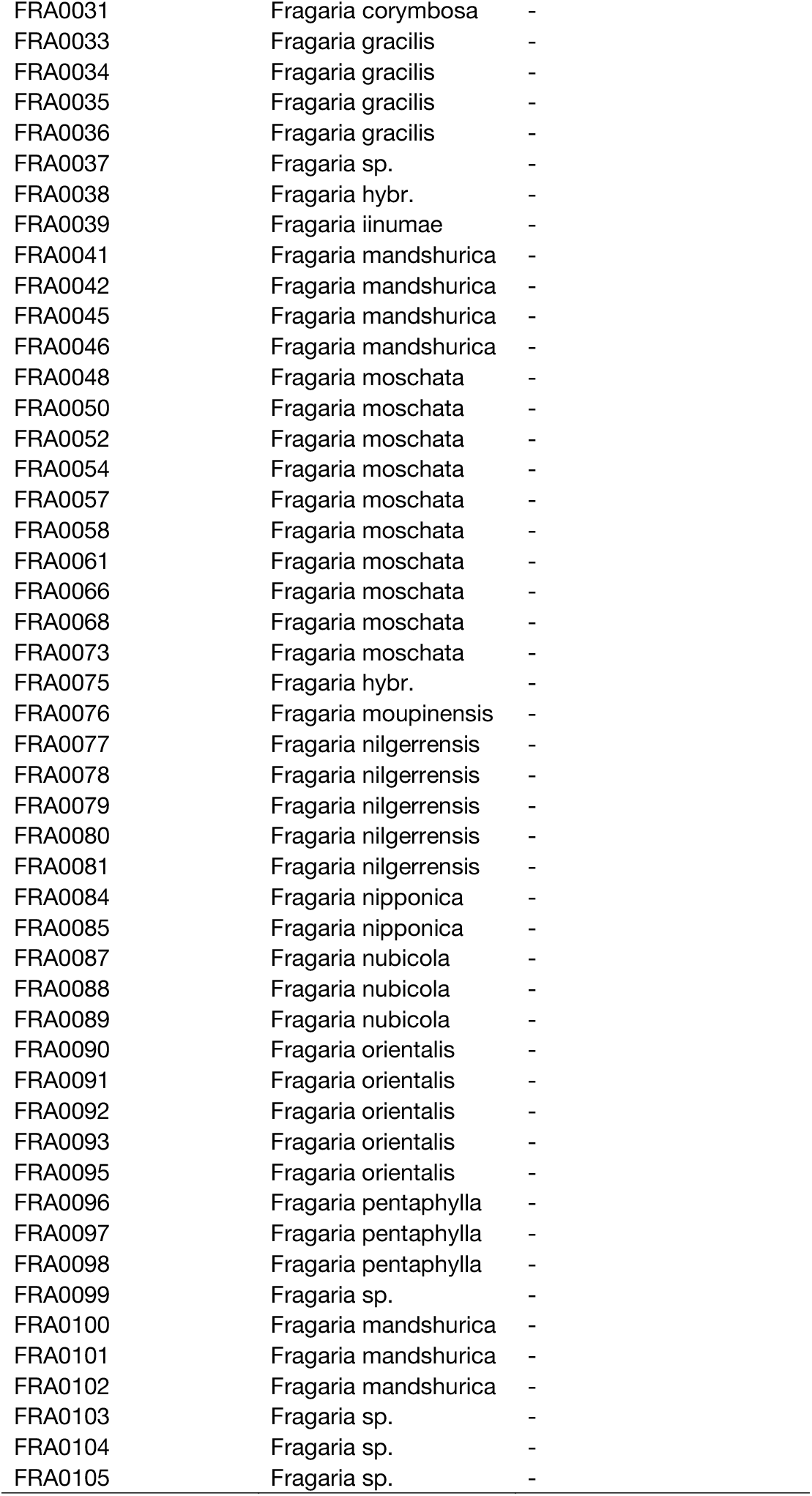
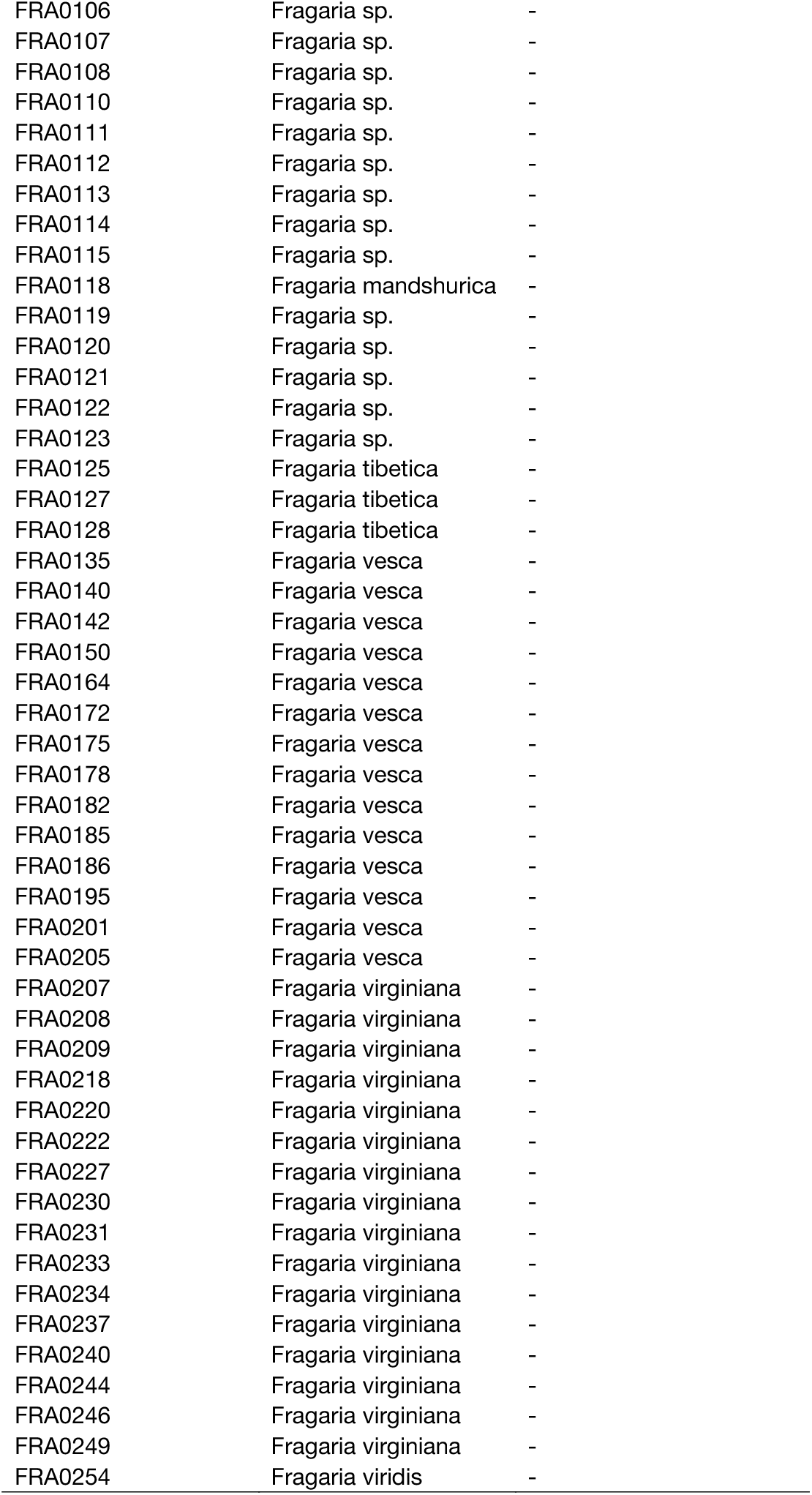
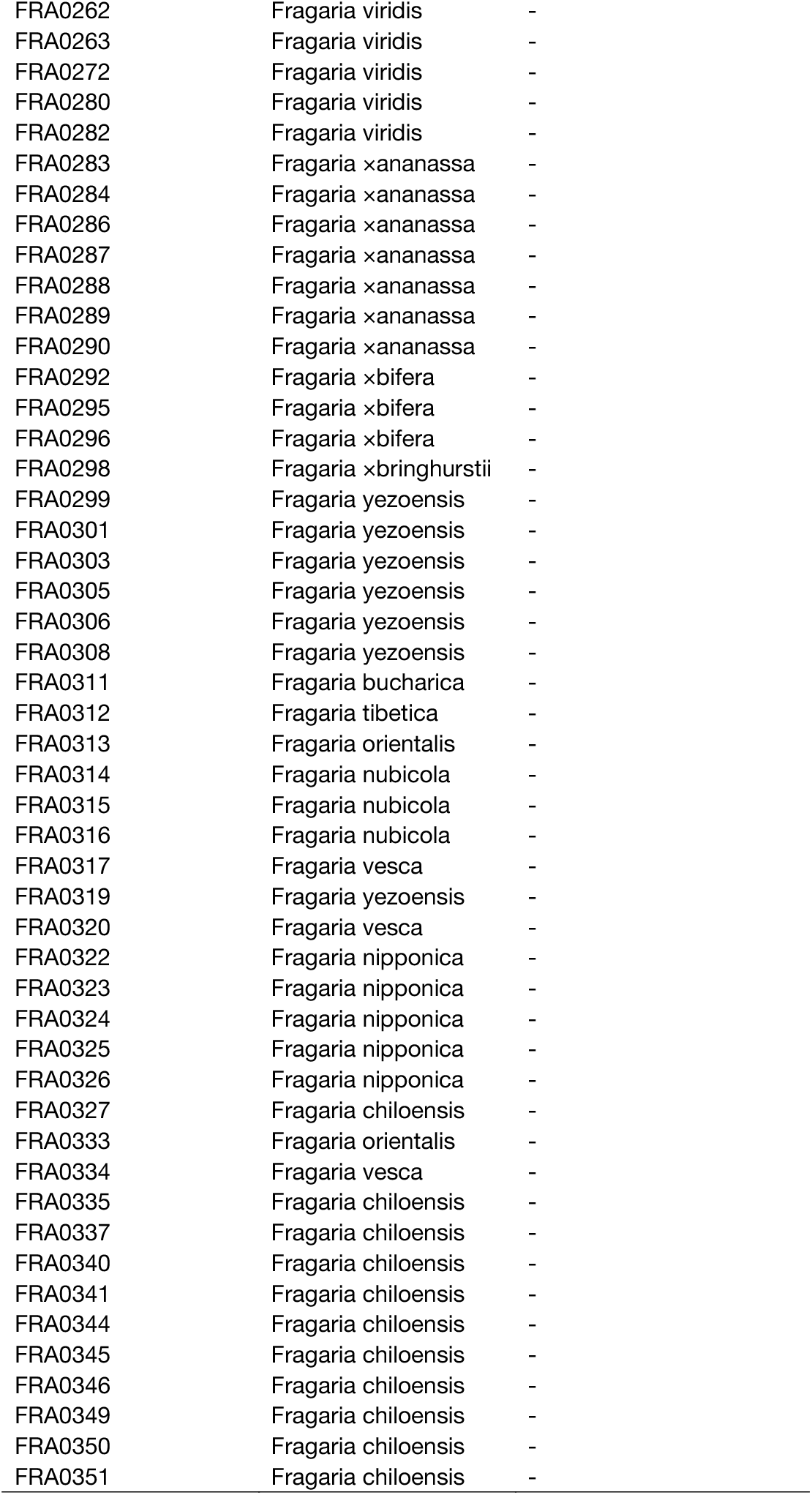
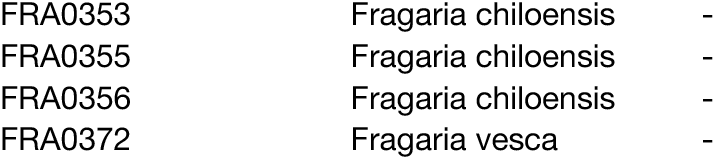
Tested accessions used in this study.

**Table S2:**
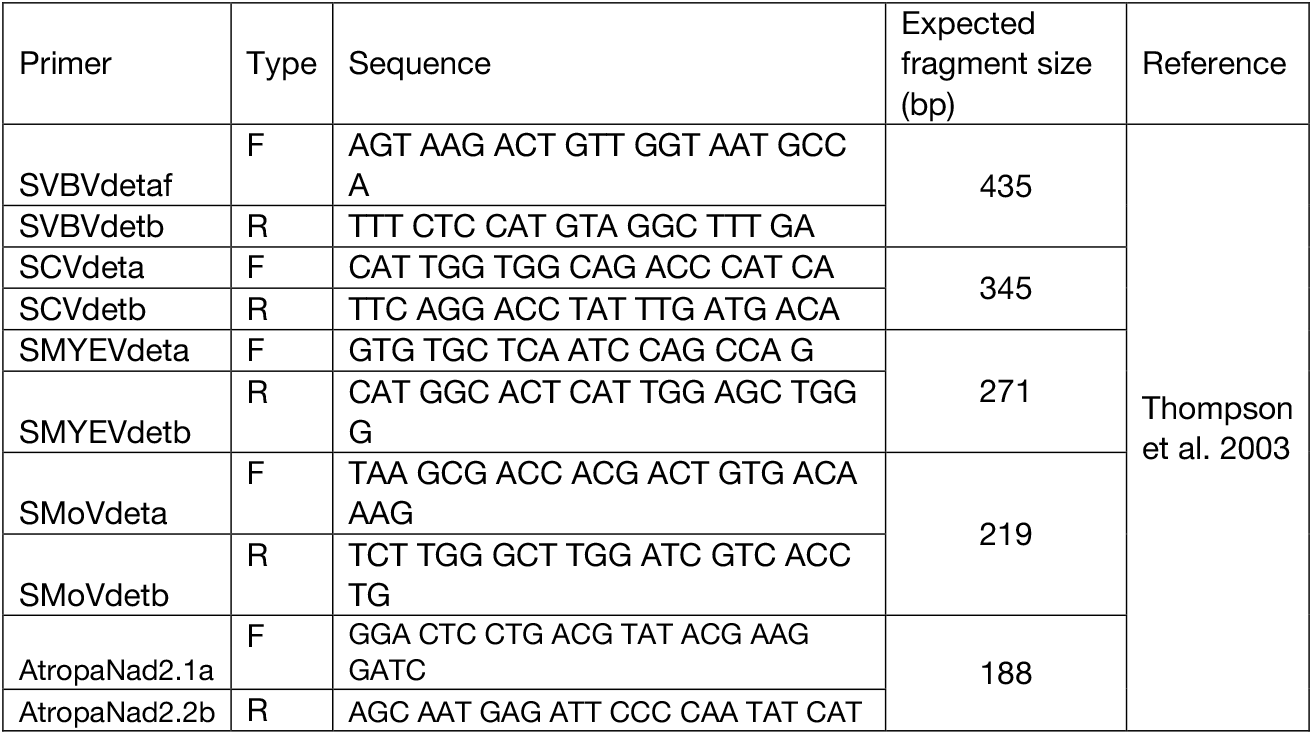
Primer sequences to proof strawberry leaf material on the occurrence of 4 strawberry viruses.

**Table S3.** Mastermix and PCR conditions for strawberry virus detection.

| Reagent (initial concentration) | µl per sample | Final concentration | PCR conditions |
| --- | --- | --- | --- |
| dd H<sub>2</sub>O | 13,4 |  | Cycler: room 215<br>Programm: SVBV<br>1 x initial denaturation: 94 °C 3'<br>38 x<br>denaturation/annealing/elongation:<br>94 C° 1'/55 °C 40''/72 °C 40''<br>1 x final elongation: 72 °C 5'<br>1 x cooling: 10 °C ∞ |
| 10 x DreamTaq Puffer (20 mM MgCl<sub>2</sub>) | 2,5 | 1 x |  |
| 2 mM dNTP's | 2,5 | 0,2 mM |  |
| SVBVdetaf (10 µM) | 1,25 | 0,5 µM |  |
| SVBVdetb (10 µM) | 1,25 | 0,5µM |  |
| 20 x rot Puffer | 1 | 0,8 x |  |
| BSA<sup>a</sup> (0,125 %) | 1 | 0,005 % |  |
| PVP<sup>b</sup> (25 %) | 1 | 1 % |  |
| DreamTaq Polymerase (5 U/µl) | 0,1 | 0,5 U |  |
| DNA-Probe<sup>c</sup> | 1 |  |  |
| Total | 25 |  |  |

<sup>a</sup> 0,005 % BSA (Bovine serum albumin)/µl was added to the mastermix to prevent PCR-inhibitory substrates (Zhang et al. 2014).
<sup>b</sup> 1 % PVP (Polyvinypyrrolidone)/µl was added to the mastermix to prevent PCR-inhibitory substrates (Koonjul et al. 1999).
<sup>c</sup> In general 10 ng/µl DNA per standard-PCR each was used, using cDNA no concentration was determined.

## Notes

### Competing Interest Statement

The authors have declared no competing interest.

